# Sex-dependent improvement in traumatic brain injury outcomes after liposomal delivery of dexamethasone in mice

**DOI:** 10.1101/2023.05.16.541045

**Authors:** Gherardo Baudo, Hannah Flinn, Morgan Holcomb, Anjana Tiwari, Sirena Soriano, Francesca Taraballi, Biana Godin, Assaf Zinger, Sonia Villapol

**Author notes:** equal contribution (Biana Godin, Assaf Zinger, Sonia Villapol).

## Abstract

Traumatic Brain Injury (TBI) can have long-lasting physical, emotional, and cognitive consequences due to the neurodegeneration caused by its robust inflammatory response. Despite advances in rehabilitation care, effective neuroprotective treatments for TBI patients are lacking. Furthermore, current drug delivery methods for TBI treatment are inefficient in targeting inflamed brain areas. To address this issue, we have developed a liposomal nanocarrier (Lipo) encapsulating dexamethasone (Dex), an agonist for the glucocorticoid receptor utilized to alleviate inflammation and swelling in various conditions. In vitro studies show that Lipo-Dex were well tolerated in human and murine neural cells. Lipo-Dex showed significant suppression of inflammatory cytokines, IL-6 and TNF-α, release after induction of neural inflammation with lipopolysaccharide. Further, the Lipo-Dex were administered to young adult male and female C57BL/6 mice immediately after a controlled cortical impact injury. Our findings demonstrate that Lipo-Dex can selectively target the injured brain, thereby reducing lesion volume, cell death, astrogliosis, the release of proinflammatory cytokines, and microglial activation compared to Lipo-treated mice in a sex-dependent manner, showing a major impact only in male mice. This highlights the importance of considering sex as a crucial variable in developing and evaluating new nano-therapies for brain injury. These results suggest that Lipo-Dex administration may effectively treat acute TBI.

---

Traumatic Brain Injury (TBI) is a significant public health concern that leads to 3 million emergency room visits annually ^1^. TBI promotes robust neuroinflammation by activating microglia and macrophages that release pro-inflammatory cytokines and chemokines. This process attracts leukocytes, neutrophils, T cells, monocytes, and infiltrating macrophages, which increase neurotoxicity and neuronal death ^2^. The invasion of immune cells contributes to a larger lesion size and a slower recovery, exacerbating neurotoxicity and neuronal death ^3, 4^. However, drug delivery to the brain has proven challenging, making pharmacotherapy for TBI ineffective. Systemic drug administration in sufficient concentration to reach the injured brain may result toxic to other organs. TBI can compromise the blood-brain barrier (BBB), which typically separates the brain from the rest of the body and regulates the passage of substances into and out of the brain. While this consequence from TBI can facilitate drug delivery to the brain, it can also allow harmful substances to enter, potentially causing further damage. Even if a drug can cross the BBB, delivering it to the specific brain region in need may be difficult. The brain is highly compartmentalized, and the drug may not be able to reach certain areas due to their location or the nature of the injury. This could potentially increase the risk of a secondary injury and disrupt the body’s natural healing processes.

Dexamethasone (Dex) is a glucocorticoid used as an anti-inflammatory agent for various conditions, including central nervous system (CNS) injury ^5^. It has been used to treat brain tumors, critical brain illness, stroke, and COVID-19 ^6–8^. Despite its efficacy, Dex cannot accumulate in the brain due to its active efflux transport by p-glycoprotein ^9^. In addition, Dex systemic administration is associated with several side effects, including glucose intolerance, immunosuppression, and neuropsychiatric problems ^9–11^. Consequently, despite a potent anti-inflammatory action, the long-term clinical outcomes of TBI patients treated with Dex are not significantly different or even worse than those treated with a placebo ^12, 13^. Thus, an efficient delivery method is needed to control Dex therapeutic concentration at the target site and limit its distribution after TBI.

Glucocorticoids, as the main physiological hormones with anti-inflammatory effect, have shown sexual dimorphism in inflammatory conditions ^14^ in inflammatory conditions. While estrogens can promote inflammation via Akt-mTOR pathway ^15^, male steroid hormones generally suppress immune function ^16^ through inhibition of NF-κB and COX, thus the effect of Dex on the inflammatory state can be sex-dependent.

Efficient drug delivery to the brain for treating neuropathological conditions remains a significant challenge. Liposomes (Lipo) are clinically used nanoparticles composed of phospholipid bilayers surrounding an aqueous core ^17^, which can improve site-specific drug delivery and shelf-life while avoiding adverse effects ^18–20^. Lipo show promise in enabling targeted delivery of drugs to the brain by circumventing efflux transporters ^21^ or specific cells targeting ^21, 22^. The compromised BBB following TBI may provide an opportunity for enhanced delivery of Lipo to damaged cortical regions. Although, Dex liposomes were used before in animal models of arthritis or *in vitro* ^23, 24^, to the best of our knowledge, and up to this date, their efficacy in a TBI model was never assessed while evaluating the sex effect on their therapeutic outcomes.

Direct delivery of anti-inflammatory agents to the affected brain areas using Lipo can potentially mitigate the harmful effects of neuroinflammation and promote recovery after TBI, while minimizing off-target effects and drug toxicity compared to other delivery methods. Our recent work shows that Lipo can be efficiently delivered to the brain after TBI ^25^. More importantly, this study also demonstrated that empty liposomes possess therapeutic benefits, as evidenced by reduced brain lesions. Nonetheless, additional research is necessary to optimize Lipo-based therapies for TBI and other neuropathological conditions, focusing on their lipid backbone and the therapeutic cargo they can deliver.

Here, by using the same liposomal phospholipid backbone with some therapeutic efficacy based on our previous work ^25^, we aimed to investigate the therapeutic potential of encapsulating Dex in it (Lipo-Dex). Our results show that Lipo-Dex reduces levels of inflammatory proteins in neuronal cell cultures treated with lipopolysaccharide (LPS) *in vitro*. In a mouse model of TBI, Lipo-Dex significantly reduced lesion volume and decreased markers of inflammation such as cell death, microglia, astrocytes, and neutrophils, demonstrating its neurorestorative effect compared to the empty Lipo. Interestingly, Lipo-Dex’s effectiveness was sex-dependent, with only male mice showing a significant reduction in inflammation and lesion size, highlighting the importance of considering sex as a factor in developing and evaluating new therapies for brain injury. Our study underscores the potential of nanotechnology for developing more targeted and effective therapies for brain injury.

## RESULTS AND DISCUSSION

### Fabrication, Characterization and Stability Assessment of Lipo and Lipo-Dex in Storage and Body Temperatures

Both Lipo and Lipo-Dex were created using the same extrusion parameters, with the exception of Dex added during lipids thin-film rehydration. The physical and chemical properties of both systems were evaluated as we previously described ^26^. Dex encapsulation had no effect on the average diameter, polydispersity index (PDI), zeta potential (ZP), and Lipo concentration (Figure 2a-c). Interestingly Lipo-Dex has a lower PDI, as compared to Lipo; but is lower than 0.2, which it is considered to be an acceptable value and indicates a homogenous population of phospholipid NPs ^27^. Lipo and Lipo-Dex showed an average size of 129.09 ± 3.449 and 115.94 ± 4.791 nm, PDI of 0.12 ± 0.0053 and 0.04 ± 0.001588 a.u., ZP of -9.98 ± 0.1655 and -9.35 ± 0.1051 mV and Lipo concentration 1.78E+13 ± 0.167 and 1.94E+13 ± 0.132 (particles/ml), respectively (N = 3). Cryo-transmission electron microscopy (cryo-TEM) images of both Lipo and Lipo-Dex (Figure 2e) demonstrated a similar bilayer structure and morphology in both formulations resulting an average size of 9.43 ± 0.12 and 8.72 ± 0.21 nm for Lipo and Lipo-Dex respectively. The thickness of the lipid bilayer was measured for 10 Lipo using the Fiji software. The release profile of Dex was assessed over 72 h at 37 °C (Figure 2i). We determined that over 80% of encapsulated Dex to be released after 4 h from Lipo-Dex according to our previous work ^28^. Furthermore, the structural stability assessment at 4 °C, followed by DLS measurements for Lipo and Lipo-Dex size, PDI, and ZP, showed no changes in the liposomal physicochemical characteristics over the 28-day period (Figure 2f-i). The Dex release profile of Lipo stored at 4 °C at the following post-synthesis points: 0, 7, 14, 21 and 28 days was evaluated. Dex release from Lipo increased by 41.6 ± 9.77%, 60.5 ± 8.75%, 71.4 ± 8.47% and 78.82 ± 7.13% at the end of 7, 14, 21 and 28 days, respectively (n = 3). The concentration of encapsulated Dex was 1.51 ± 0.14 mg/mL (n = 3) with 51.2% Dex entrapment capacity (n = 3). The release of Dex from Lipo stored at 4 °C increased over time, and the reproducibility of the synthesis process was confirmed. These results are consistent with previous findings for lipid-based formulations ^29^. These results suggest the successful fabrication of Lipo and Lipo-Dex systems using the same extrusion parameters with no significant differences in their physiochemical properties. Previous studies have demonstrated the importance of consistent physiochemical properties in lipid nanoparticle formulations. For example, as reviewed by Hoshyar et al. (2016) variations in the size and surface charge of lipid nanoparticles can significantly affect their cellular uptake and efficacy ^30^. Similarly, it was demonstrated that the stability of lipid nanoparticles can be influenced by their physiochemical properties, which in turn can affect drug release and cellular uptake ^31^. The release of Dex from Lipo-Dex in our study is consistent with previous findings for lipid-based formulations, as reported in studies on the release of other corticosteroids such as methylprednisolone (Xiao et al. 2018) and hydrocortisone ^32^.

**Figure 1.**
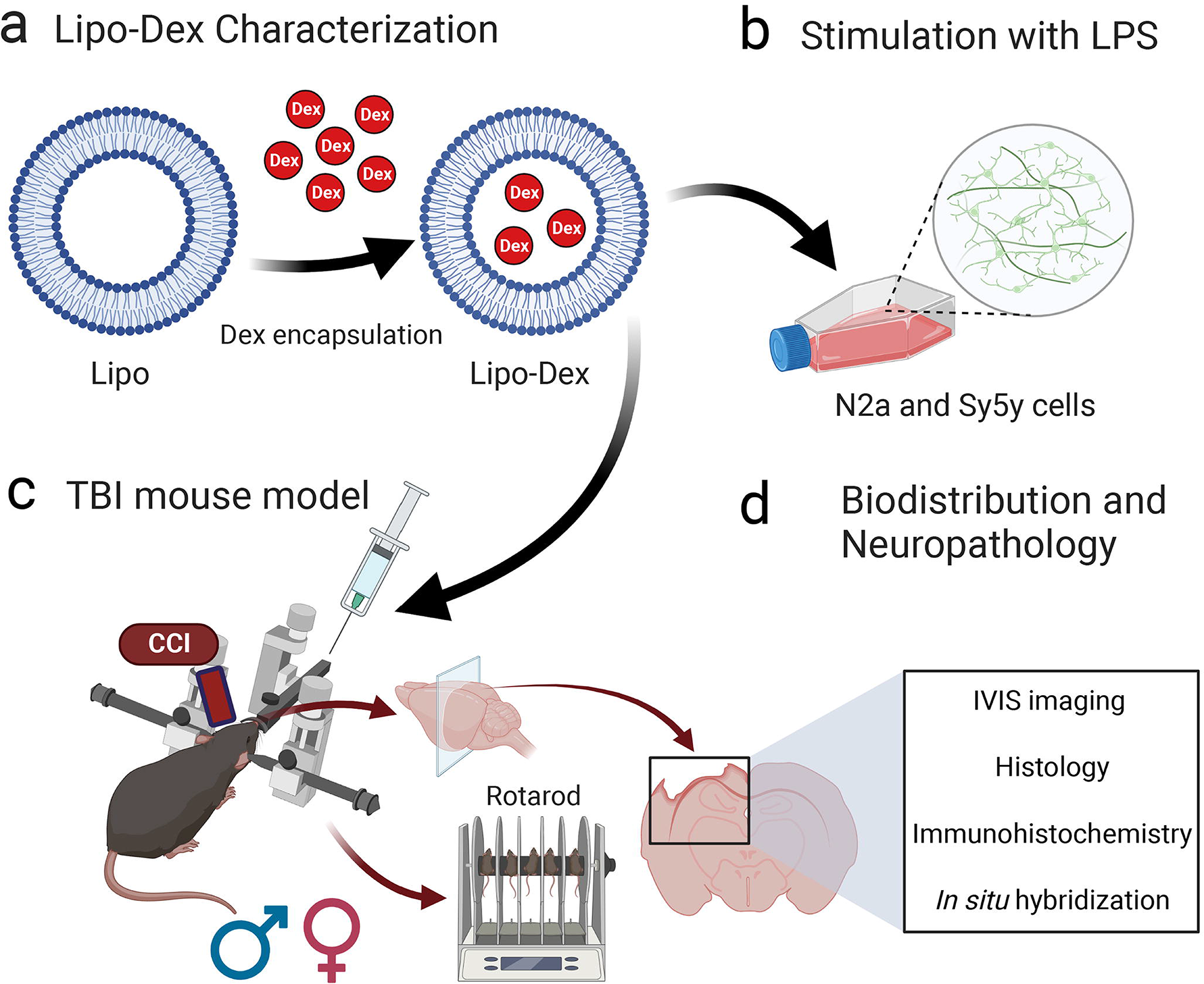
Schematic of Lipo-Dex fabrication, characterization, and *in vivo* and *ex vivo* experiments. (a) Dex encapsulation was performed *in vitro* and after fabrication, Lipo-Dex were characterized for their physicochemical properties and Dex release measurements. (b) neuronal cells (N2a and Sy5y) were stimulated with LPS and inflammatory cytokines were measured. (c) Lipo-Dex were tested *in vivo* using a traumatic brain injury (TBI) mouse model, called controlled cortical impact (CCI) injury, and (d) the therapeutic effect was evaluated using *in vivo* imaging system (IVIS), histological, immunohistochemical, and *in situ* hybridization techniques while assessing the mice sex effect on the therapeutic outcomes.

**Figure 2.**
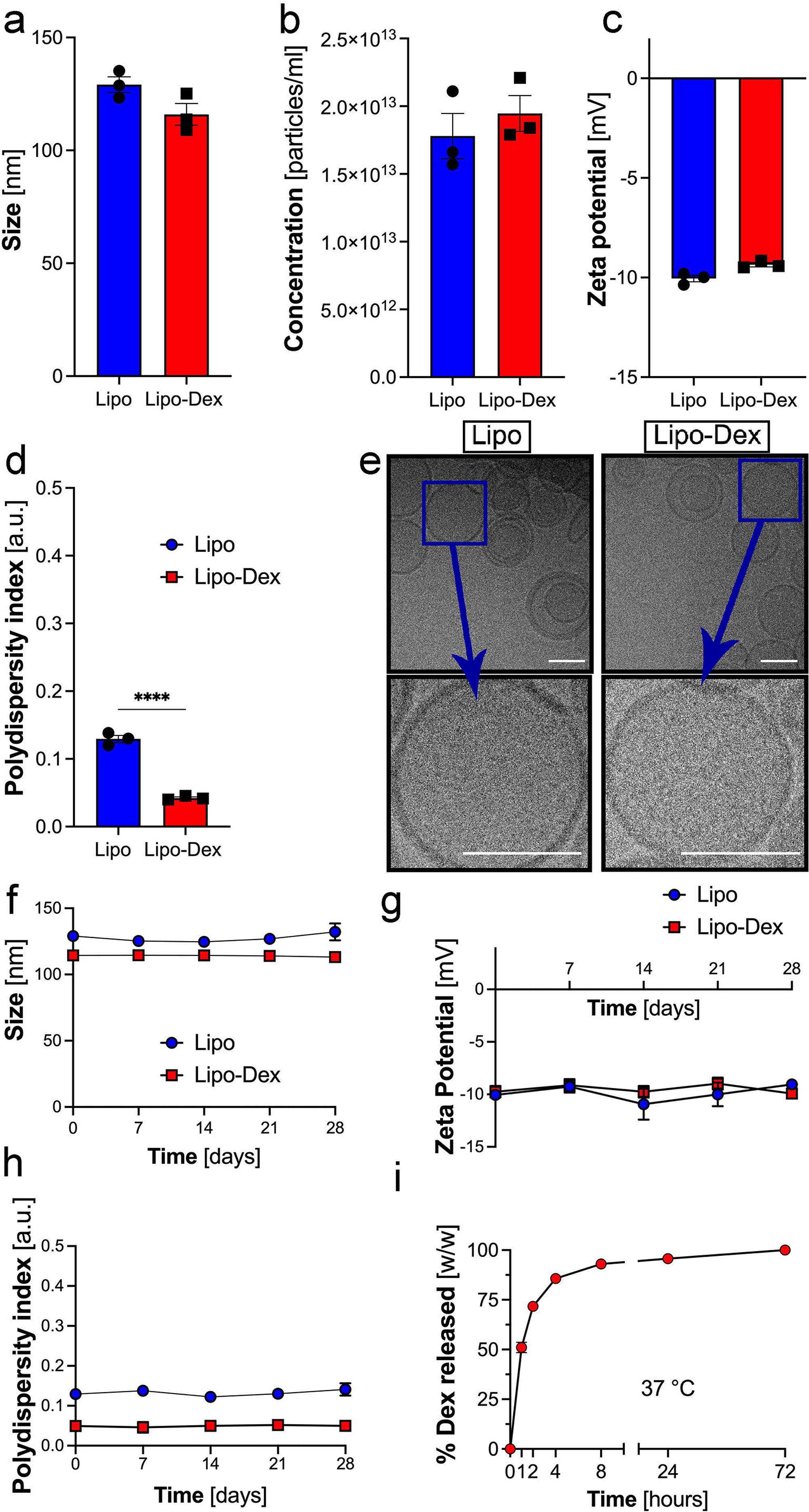
Fabrication, Characterization and Stability Assessment of Lipo and Lipo-Dex in storage and body temperatures. Lipo and Lipo-Dex were characterized for their physicochemical properties using DLS. No significant changes in (a) size, (b) concentration and (c) zeta potential (ZP) were observed. However, while both systems show PDI < 0.2, indicating a very homogeneous nanoparticle population (d), Lipo-Dex have a lower PDI, as compared to Lipo. (e) Representative cryo-TEM images of Lipo and Lipo-Dex verified that no structural changes occurred after encapsulating Dex within Lipo. Scale bar represents 100 nm for all the images. Lipo and Lipo-Dex were stable in terms of (f) size, (g) ZP, (h) PDI over a period 28 days stored at 4 °C. (i) Lipo-Dex show a burst Dex release (∼80%) after 4 h at 37 °C. Results are shown as mean ± SEM. ****p<0.0001.

### Biocompatibility and Efficacy of Lipo-Dex Suppressing the Inflammation in Neuronal Cells

We first evaluated the biocompatibility of Lipo-Dex *in vitro* using WST-1 assay in Sy5y human neuronal cells and N2a murine neural cells and did not Lipo-Dex were well tolerated over Dex concentration range of 0.001 to 10 µM in both cell lines, and did not show a significant difference in the cell viability compared to the untreated control at 24 and 48 h post-treatment (Figure 3a, b). Further, we tested the ability of Lipo-Dex to suppress LPS-induced inflammation *in vitro*. When administered at a concentration of 5 µg/ml, LPS significantly increased the expression levels of inflammatory cytokines IL-6 and TNF-α in Sy5y cells compared to those administrated with no LPS. This inflammatory response has been previously reported ^33, 34^. Lipo-Dex at a Dex concentration of 1 µM and 5 µM significantly suppressed the release of IL-6 and TNF-α (Figure 3 c-d). IL-6 and TNF-α are the major inflammatory factors in the CNS that mediate inflammation related to trauma ^35, 36^. Previous studies have shown that Dex can prevent LPS-induced inflammatory responses in neuronal cells ^37^. Interestingly, a study with a different liposomal formulation of Dex demonstrated a reduction in TNF-α and IL-6 secretion in inflamed primary human macrophages ^24^. Our findings show that while being biocompatible in non-LPS treated cells, Lipo-Dex were able to mitigate LPS-induced inflammation.

**Figure 3.**
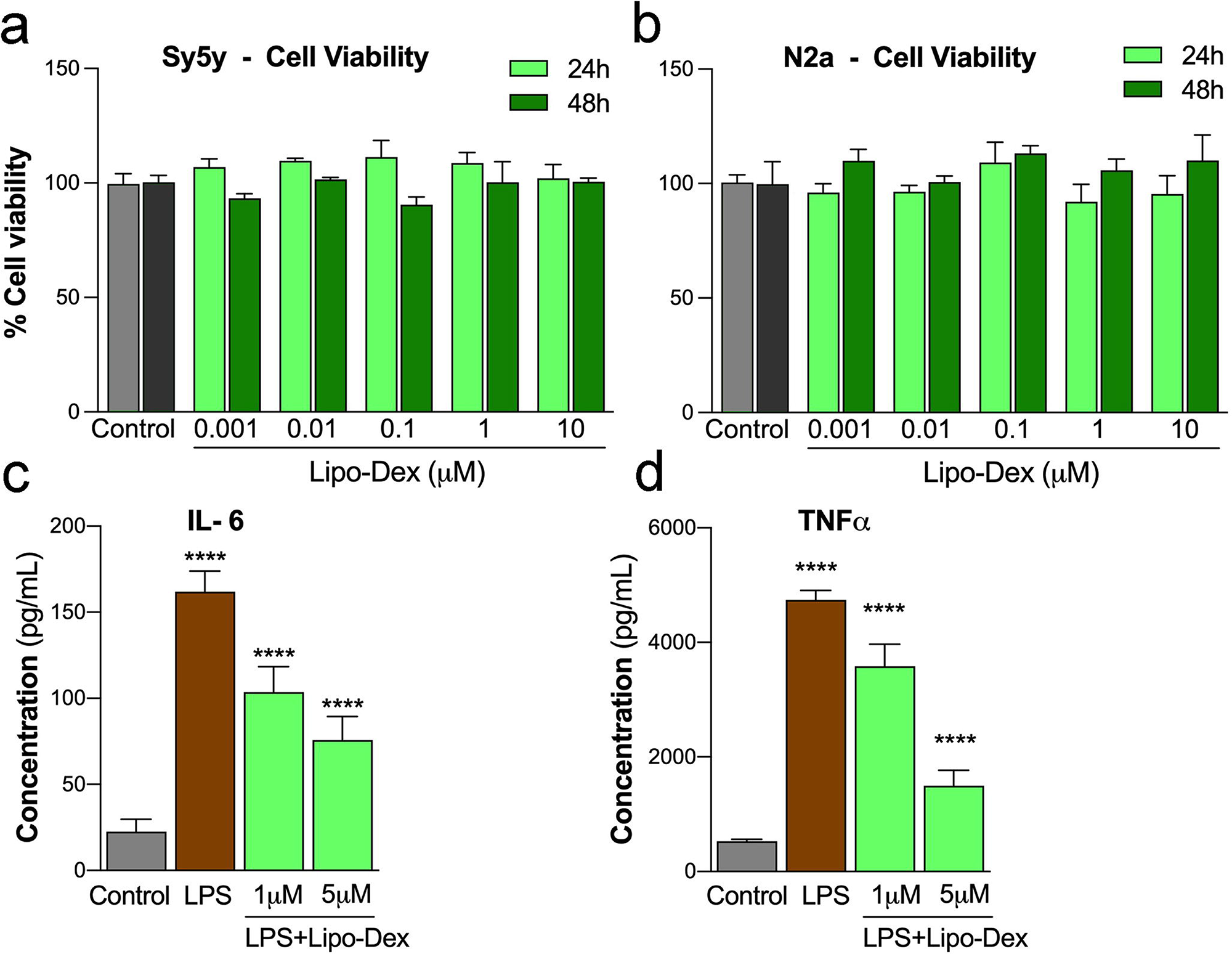
Lipo-Dex is biocompatible and decreases inflammation in neuronal cells *in vitro.* *In vitro* biocompatibility of Lipo-Dex in neuronal cells. Sy5y human neuronal cells (a) and N2a mouse neuronal cells (b) incubated with different concentrations of Dex delivered by Lipo-Dex. Sy5y neuronal cell line pre-treated with Dex (5µM) delivered from Lipo-Dex for 1 h before induction of inflammation with LPS (5 µg/ml) for 24 h. IL-6 (c) and TNF-α (d) levels were reduced in Sy5y cells inflamed with LPS and then treated with Lipo-Dex for 24 h. Data are expressed as mean ± SEM from three independent experiments. ***p<0.001. No significant differences were found between Lipo-Dex treated and untreated cells control groups, indicating the biocompatibility of the system with neuronal cells.

### Lipo-Dex Bioistribution and Organ Accumulation in a Mouse Model of TBI

Next, we set out to characterize the effect of Lipo-Dex *in vivo* following TBI using a mouse model of moderate TBI, well established in our laboratory ^38, 39^, followed by intravenous administration of Lipo-Dex 10 min after injury. We evaluated the biodistribution of Lipo-Dex by using Cy5.5-labeled Lipo and *in vivo* imaging with IVIS. We found that Lipo-Dex were present in several organs of the TBI mice (Figure 4). Lipo-Dex demonstrated significant accumulation in the brain at the site of the lesion (Figure 4), consistent with our previously reported data on the biodistribution of empty Lipo in TBI mice ^25^. Lipo-Dex were also accumulated in the filtering organs, spleen and liver, lungs and kidneys, but was not detected in the heart and blood. Male and female mice in the Lipo-Dex group exhibited similar patterns of Lipo accumulation in the brain lesions and other tissues except for spleen, showing a significant increase in Lipo accumulation in male mice compared to female mice (Figure 4). Subsequently, to evaluate if both Lipo-Dex and Lipo were tolerated *in vivo*, the mentioned organs were collected, washed, fixed, sectioned, and stained using hematoxylin and eosin (H&E) to evaluate tissue damage. When compared with tissue from a control PBS-injected mouse, no abnormal or pathological morphology differences were observed in any of the two groups’ organs (Figure S1). In our previous study where we administered Lipo and leukosomes after TBI ^25^, there were no significant sex differences in the biodistribution of Lipo-Dex. We also found an accumulation in the brain, spleen and liver. However, in the present study, we noticed a lower accumulation of Lipo-Dex in the lungs, heart, and kidneys overall. This implies that despite using a systemic administration, the Lipo are directed mainly to the inflamed brain.

**Figure 4.**
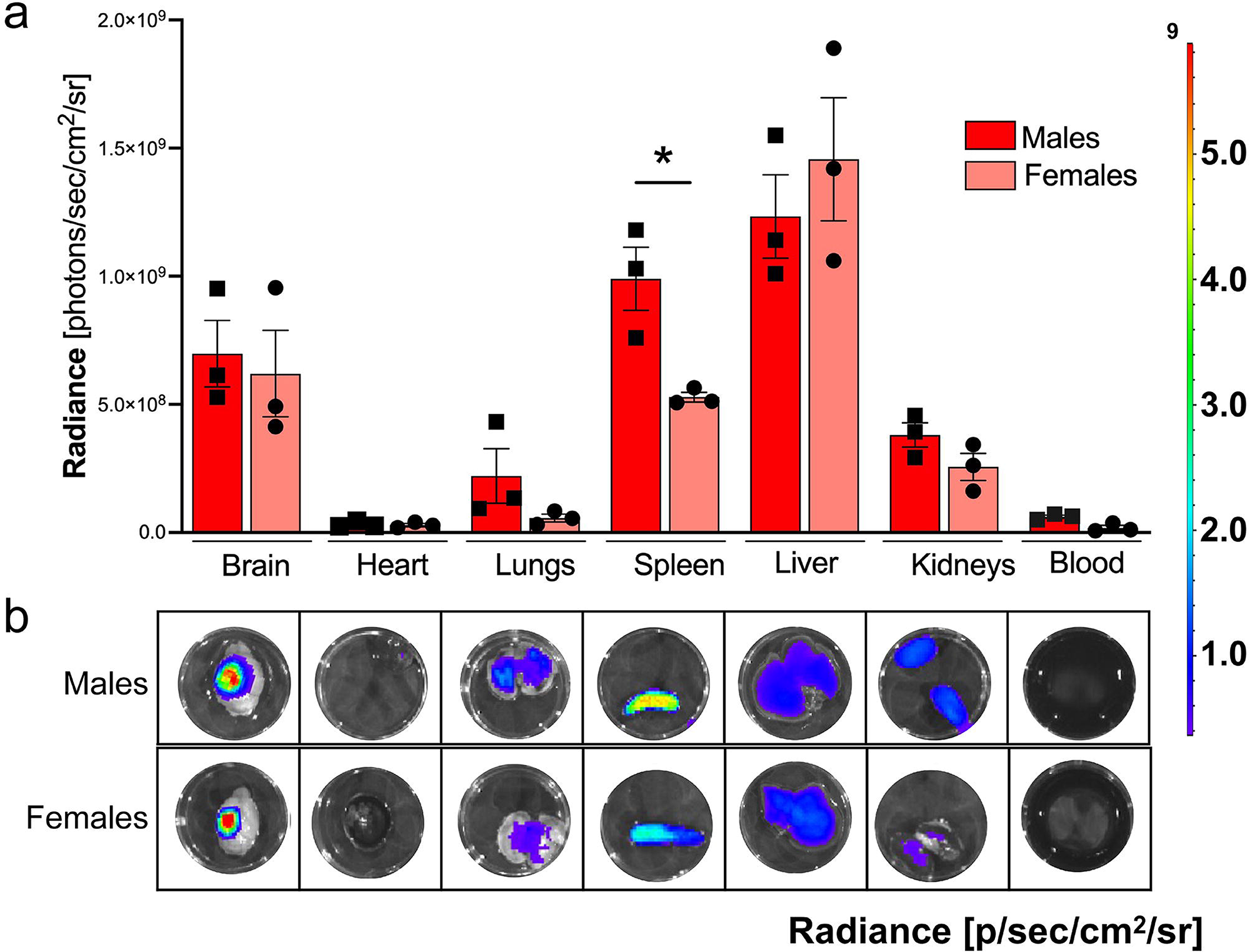
Sex dependent *lipo-Dex biodistribution and brain targeting in vivo in traumatic brain injury (TBI) mouse model.* Both male and female mice were administrated Cy5.5-labeled Lipo-Dex via retro-orbital injection under anesthesia. (a) Lipo-Dex demonstrated higher accumulation in brain and filtering organs (spleen and liver). Interestingly, there was a significant higher accumulation in the spleens of males compared to females. One-way ANOVA followed by Tukey’s multiple comparison test was used to determine statistical probabilities between brain biodistribution of Lipo-Dex in male and female mice for heart, lungs, spleen, liver and kidneys and blood. Results are shown as mean ± SEM. \**p* ≤ 0.05, *n* = 3/group. (b) Representative image of each organ (brain, heart, lungs, spleen, liver, and kidneys) and blood.

### Neuroprotective Effects of Lipo-Dex *in vivo* in TBI Mouse Model

To assess the acute neuroprotective effects of Lipo-Dex after TBI, we quantified cell death in the pericontusional area and the lesion volume. The findings showed a significant reduction of 54% in Terminal deoxynucleotidyl transferase dUTP nick end labeling (TUNEL)-positive cells (Figure 5a-e) and a reduction of 38% in lesion volume (Figure 5f) in male mice treated with Lipo-Dex compared to those treated with Lipo. Interestingly, no significant differences were observed between the two groups in females. Administration of Lipo-Dex through microcirculation may be a suitable method for delivering the drug to the brain in effective doses. Our previous study found that empty Lipo effectively reached injured and inflamed brain areas ^25^. In the present study, we found that encapsulating Dex in Lipo reduces the inflammatory response and reduces the lesion volume in a TBI mouse model. Our study also revealed that sex affects the efficacy of Lipo-Dex treatment, with only male mice showing a significant reduction in inflammation and lesion size. There were higher endogenous antioxidant enzyme activities in the brains of female than in male mice ^40, 41^, thus the potential antioxidant effect of Dex works only in males in the acute TBI response. These findings emphasize the importance of considering sex when developing and evaluating new therapies for TBI. To evaluate the motor ability after Lipo-Dex administration in TBI mice, a rotarod test was conducted 24 h post-injury. There was a reduction in motor ability in all injured animals compared to baseline values, as expected. The findings indicated that there was no significant improvement in functional motor ability in either male or female mice treated with Lipo-Dex compared to those treated with Lipo (Figure 5g). Other studies have shown that Dex exhibited significantly improved motor functions and cognitive behavior compared to untreated mice at 14 days post-TBI ^42^. We have to highlight that the present study focuses on characterizing the effect of Lipo-Dex in the acute stage after TBI and treatment differences in motor function may be taking place in later timepoints.

**Figure 5.**
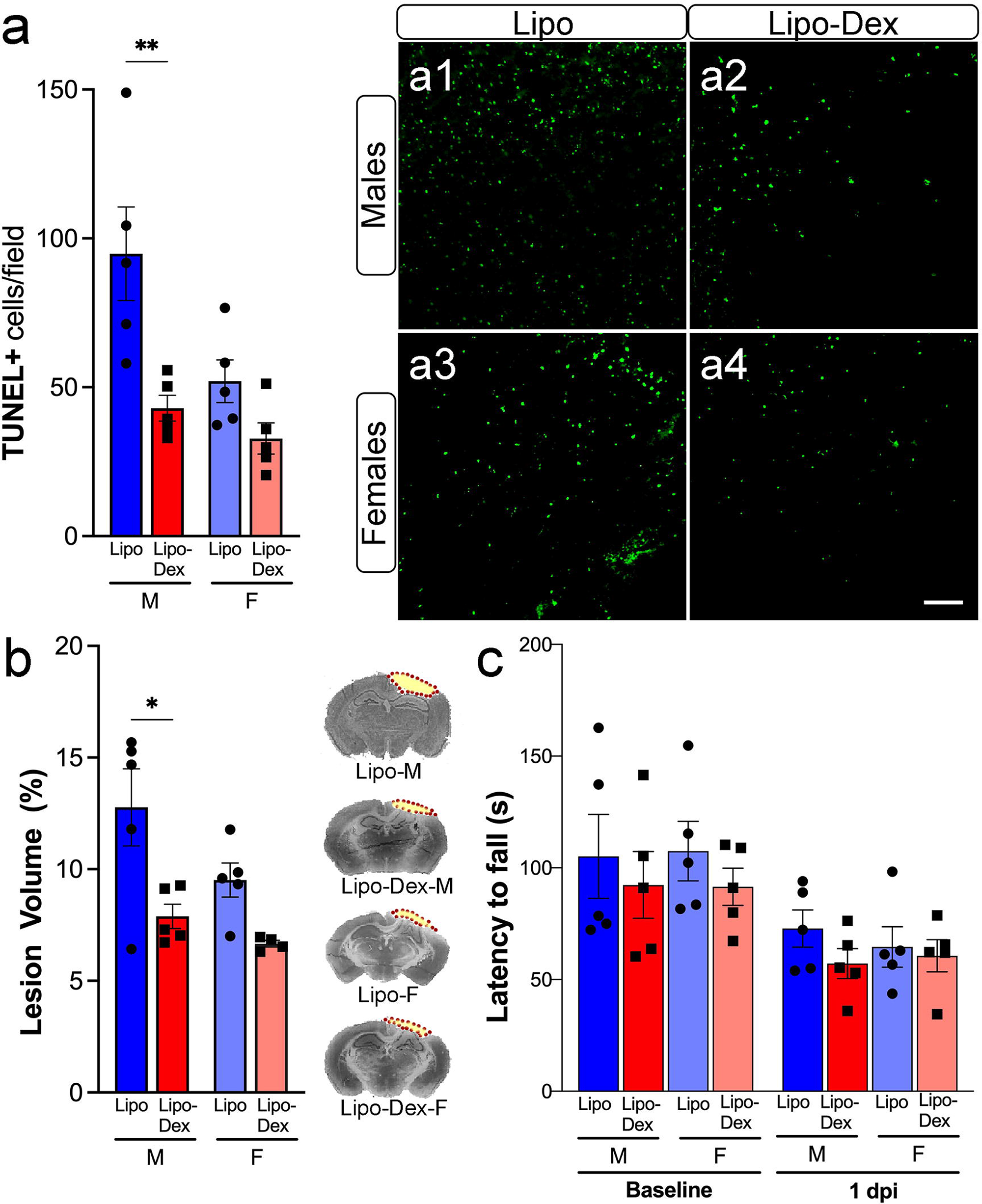
Lipo-Dex treatment reduces the apoptotic cells and lesion volume in the injured cortex but did not improve the motor ability after TBI. Lipo-Dex administration reduces the number of TUNEL positive cells (a, a1-a4) and lesion volume (b) in male mice at 1-day post-TBI. Representative images of brain sections stained with cresyl-violet 24 h after TBI. The dotted line indicates the lesion area composed of the cavity and edematous area (b, right). Scale bar in a1-4 represents 50 μm. Lipo-Dex-treated mice did not improve motor recovery using the rotarod (c). Results are shown as mean ± SEM with \**p* ≤ 0.05, \*\**p* ≤ 0.01, *n* = 5/group.

### Lipo-Dex Reduces the Microglia and Macrophage Density and Pro-Inflammatory Cytokines in the Brain, but not in the Peripheral Blood Following TBI

We evaluated the effect of Lipo-Dex on the acute inflammatory response characteristic of TBI brains, including microglia/macrophage infiltration and release of pro-inflammatory cytokines. For this quantitative analysis, we focused on the primary somatosensory cortex area adjacent to the impact region surrounding the injury site. Following TBI, there was a rapid increase in the number and percentage of Iba-1 positive cells in the injured cortex and hippocampus, which was significantly reduced in the Lipo-Dex group at 24 h post-TBI in male mice but not in females (Figure 6a, a1-4, and b, b1-b4). The administration of Dex intravenously after TBI has been previously shown to reduce the activation of activated microglia at 14 days post-TBI ^43^. Also, differences between male and female mice suggest important sex-dimorphism responses to antioxidant nanoparticle delivery and that there may exist a maximal benefit from local antioxidant activity in injured brain ^44^. It was defined that female mice had higher endogenous antioxidant activity, which delayed the increase in post-traumatic markers of oxidative stress after TBI compared to male mice. We further investigated the pro-inflammatory response by measuring the mRNA expression of the cytokines TNF-α and IL1-β in injured brain sections using fluorescent *in situ* hybridization (FISH). We observed a trend towards a reduced IL1-β (Figure 6c, c1/c4) and TNF-α (Figure 6d, d1-d6) expression in the ipsilateral peri-contusion region in Lipo-Dex-treated males compared to Lipo-treated males, but not in females. Other studies showed that Dex-induced TNF-α overexpression may affect astrocytic hypertrophy without affecting microgliosis, indicating a possible role in neuronal function after TBI ^45^. There was no detectable IL1-β and TNF-α mRNA expression in the contralateral hemisphere (data not shown). TNF-α and IL1-β mRNA expression combined with Iba-1 antibody using FISH demonstrated that most of the cytokine signal colocalized with microglia/macrophages markers (Figure 6 e1-4, f1-4). We also assessed the changes in pro-inflammatory and anti-inflammatory cytokines in the plasma to determine if there is a systemic anti-inflammatory effect following Lipo-Dex administration. Dex administrated intraperitoneally 1 h post-TBI significantly reduced the expression of proinflammatory cytokines such as IL1-β, TNF-α, and IL-6 in the brain, indicating its potential as an anti-inflammatory agent ^46^. Dex has also been shown to reduce the expression of immune response genes, including MCP-1/CCL2 and ICAM-1, which return to control levels ^47^. Furthermore, Dex regulates NF-KappaB levels, which increase in brain tissue early after TBI and induce an inflammatory response that can cause secondary brain injury. By preventing this secondary injury caused by inflammatory cytokines, Dex appears to have a protective effect on the injured brain ^48^. Our data showed no differences in serum concentrations of IL1α, IL-6, IL-3, or IL-10 between Lipo and Lipo-Dex-treated mice, in either in males, nor females, at 24 h post-TBI (Figure 6 g-j). This indicates that the anti-inflammatory effect of Dex was localized in the brain injured regions without having an effect on blood peripheral inflammation.

**Figure 6.**
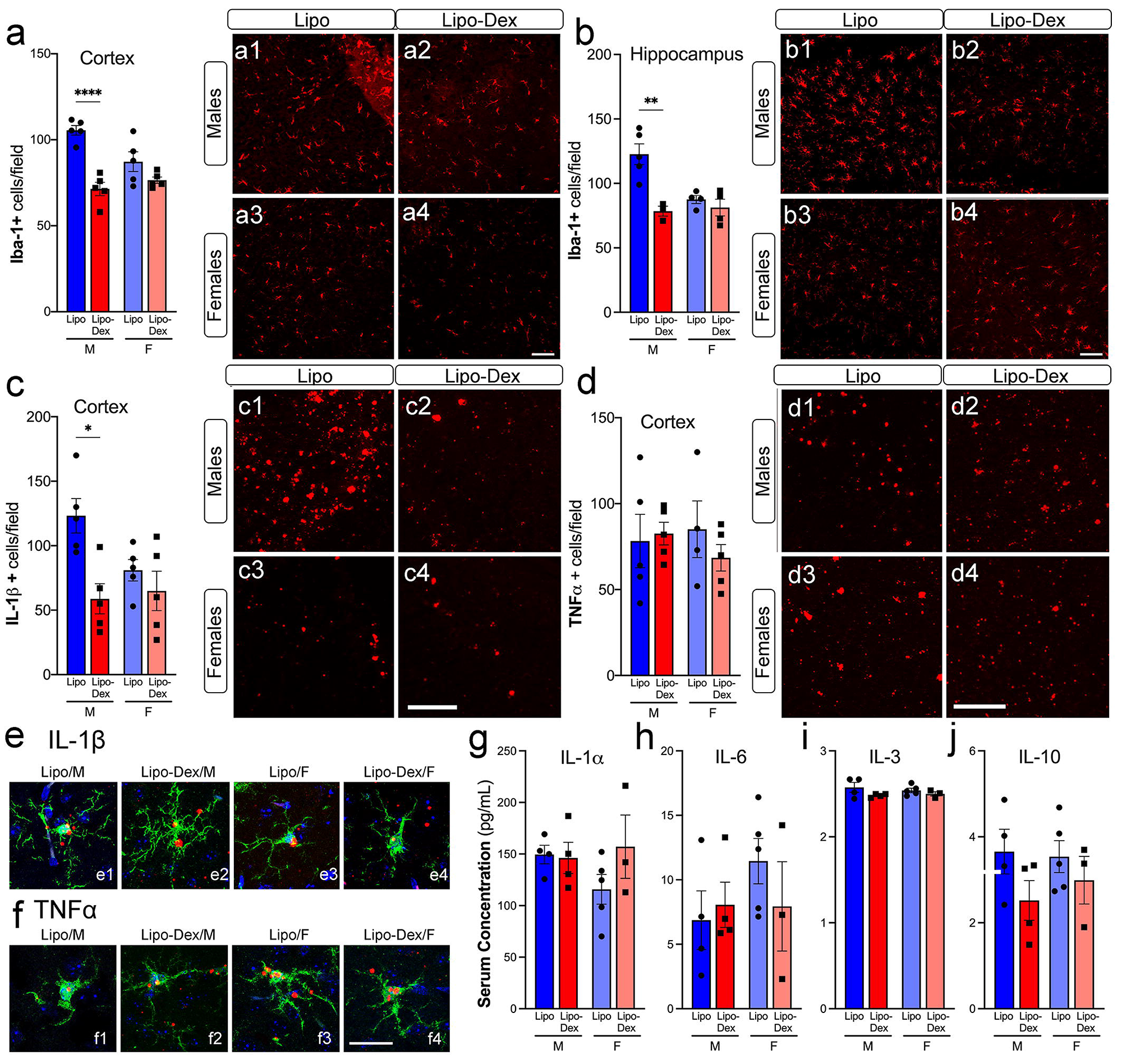
Lipo-Dex reduces neuroinflammation following TBI. Microglia/macrophages (Iba-1, red) positive cells decreased in males after TBI in the cortex (a, a1-a4) and hippocampus (b, b1-b4) compared to Lipo-treated mice. The density of IL-1b mRNA expression in the cortex (c, c1-4) was reduced after Lipo-Dex treatment, however we did not find changes in the TNF-α mRNA expression (d, d1-4). High-magnification confocal images show the colocalization between Iba-1 marker and IL1-β (e) and TNF-α (f) mRNA cytokine expression. The levels of pro-inflammatory cytokines IL-1 α (g), IL-6 (h), IL-3 (i), and anti-inflammatory IL-10 (j) are not altered after Lipo-Dex treatment in serum blood samples. Scale bar represents 50 μm for a1-4, b1-4, c1-4, d1-4, and 20 μm for e1-4 and f1-4. Results are shown as mean ± SEM. \*\**p* < 0.01; \*\*\*\**p* < 0.0001, *n* = 5/group.

### Lipo-Dex Treatment Reduces Astrogliosis Following TBI

To determine whether Lipo-Dex treatment altered the activation or survival of astroglial cells, we examined staining for glial-specific markers in brain sections taken from Lipo or Lipo-Dex treated mice at 1-day post-TBI. We used GFAP and S100β immunohistochemistry to assess the temporal and spatial distribution of astrogliosis and astrocyte density in male and female brains following TBI. Lipo-Dex treatment significantly decreased the number of S100β-positive cells and the amount of GFAP immunoreactivity in males and female brains at 1-day post-TBI (Figure 7). Lipo-Dex administration also led to a change in the morphology of the astrocytes in the cortex (Figure 7 a, b, b1-4) and reduced the number of astrocytes that were S100β-positive in the cortex and corpus callosum at 1-day post-TBI (Figure 7 c, d, d1-4). While some studies have failed to demonstrate that Dex can prevent or ameliorate gliosis in a neurodegenerative mouse model ^49^, others have found positive effects of Dex in different disease models. For example, in an epilepsy model, Dex administration has been found to improve inflammation, prevent astrogliosis (based on GFAP and S100B expression), and partially reduce astroglial dysfunction ^50^. Dex significantly reduced the GFAP signal in transgenic mice that overexpress GFAP, as measured by bioluminescent imaging ^51^, and caused a significant reduction in the number and density of astrocytes in the hippocampus and corpus callosum of postnatal rats ^52^.

**Figure 7.**
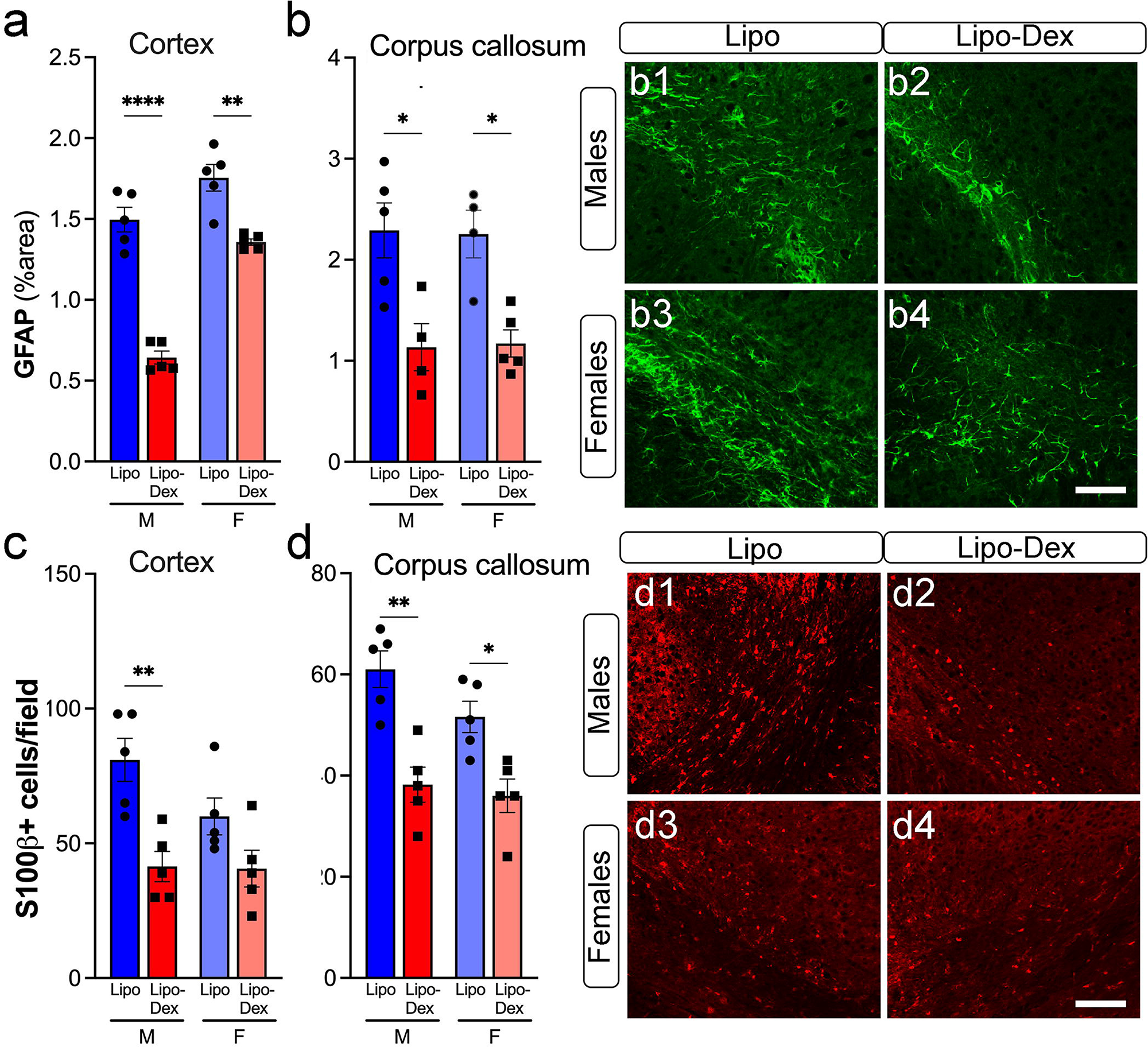
Lipo-Dex treatment reduces astrogliosis after TBI. Astrogliosis (GFAP positive cells, green) decreases after Dex treatment in the cortex (a) and corpus callosum (b, b1-b4) in both males and females at 1-day post-TBI. Astrocytic density marked with S100b was decreased in Lipo-Dex-treated mice compared to Lipo-treated after TBI in the cortex (c) in males and in the corpus callosum in males and females (d, d1-d4). Two-way ANOVA followed by multiple comparison test were used to determine statistical probabilities. *n* = 5/group. Scale bar represents 50 μm. Results are shown as mean ± SEM. \**p* < 0.05; \*\**p* < 0.01; \*\*\*\**p* < 0.0001.

## CONCLUSIONS

Our study reveals that encapsulating Dex in Lipo presents a promising approach for effective glucocorticoid delivery to young adult male and female C57BL/6 mice immediately following TBI, with more pronounced efficacy in males. This approach leverages the small size of Lipo, enabling them to access the brain vasculature through the bloodstream, which is particularly advantageous when the BBB is compromised by TBI. Our results showed that Lipo-Dex accumulated in the injured brain, resulting in reduced lesion volume, cell death, astrogliosis, release of proinflammatory cytokines, and microglial activation compared to mice treated with Lipo alone. In addition, our *in vitro* studies demonstrated the favorable tolerance of Lipo-Dex in human and murine neural cells. Furthermore, Lipo-Dex exhibited significant suppression of inflammatory cytokines, IL-6 and TNF-α, after inducing neural inflammation with lipopolysaccharide. Interestingly, these effects were found to be sex-dependent, with a significant impact observed only in male mice, highlighting the importance of considering sex differences when administering this therapy, as male and female patients may exhibit differing responses to treatment. Interestingly, these effects were found to be sex-dependent, with a significant impact observed only in male mice. Taken together, our findings suggest that drug delivery methods, such as encapsulating drugs in Lipo, while also utilizing the effect of the carrier itself ^25^ may offer new clinical opportunities for treating TBI and other neuropathological conditions. However, more research focusing on different lipids compositions and various liposomes physiochemical properties is necessary to optimize this approach better understand the mechanisms underlying the observed sex-dependent responses. Further research and clinical trials are warranted to explore the translational potential of Lipo-Dex in human TBI patients and to fully elucidate the effects of Dex, and the delivery system used to deliver it for improving brain function and recovery after TBI.

## MATERIALS AND METHODS

### Lipo-Dex Design and Purification

To encapsulate Dex in Lipo, we followed our previously published protocol ^25^ (Figure 1a). Initially, Lipo and Lipo-Dex were designed using a 4:3:3 molar ratio of Dioleoylphosphatidylcholine (DOPC), dipalmitoylphosphatidylcholine (DPPC), and cholesterol (Avanti Polar Lipids, Inc., Alabaster, United States) at a lipid concentration of 50 mM. Next, the lipids were dissolved in chloroform (Sigma Aldrich, Missouri, US) at a final concentration of 20 mg/mL, mixed, and sonicated for 5 min at 45 °C. The chloroform was then evaporated using a rotary evaporator (BUCHI Labortechnik AG, Flawil, Switzerland) at 45 °C, 280 rpm for 30 min, and the lipid film obtained was hydrated for 30 min at 50 °C and 280 rpm with either 2 mL phosphate-buffered saline (PBS, Fisher Scientific, Hampton, US) for Lipo or 2 mL of 25 mg/mL dexamethasone (Sigma-Aldrich, St. Louis, US) for Lipo-Dex. Subsequently, Lipo and Lipo-Dex were extruded at 50 °C through 80-200 nm polycarbonate membranes (Whatman, Buckinghamshire, UK). For biodistribution studies, 0.1 mg of Cy5.5-DSPE (Avanti Polar Lipids, Inc, Alabaster, US) were dissolved in chloroform (100 µL of 1 mg/mL) together with other lipids to fluorescently label both Lipo groups. Lipo-Dex were dialyzed for 19 h in PBS using 1000 kDa float A-Lyzer (Spectrum™ Labs, Massachusetts, US), with the buffer exchanged after 1 and 3 h. Subsequently, the samples were sterilized using a 0.22 µm PVDF filter.

### Lipo and Lipo-Dex Characterization

Liposomal systems were characterized using dynamic light scattering (DLS) (Malvern Instruments, Worcestershire, UK) to determine their diameter, polydispersity index (PDI), and zeta potential (ZP). Three measurements, consisting of 10 runs each, were taken to determine the size and PDI of the liposomes, while ZP was measured using three measurements, consisting of 15 runs each. The fluorescence of Lipo-Dex labeled with Cy5.5 lipid was assessed at 678 nm excitation and 707 nm emission using the FLUOstar Omega microplate reader (BMG, Labtech Ortenberg, Germany).

### Encapsulation and Drug Release Studies

The efficiency of Dex encapsulation in Lipo and its subsequent release was evaluated with high-performance liquid chromatography (HPLC) using the 1260 Infinity II LC System (Agilent Technologies, California, US). A Luna^®^ 5 μm C18(2) 100 Å, LC Column 250 x 4.6 mm (Phenomenex, California, United States) was used. The samples were mixed with PBS and acetonitrile (ACN) at a ratio of 1:6:3 (v/v/v) and sonicated, followed by centrifugation and transfer to HPLC vials. The tubes were warmed for 10 min at 40 °C and then sonicated for 10 min at 40 °C. Subsequently, the samples were transferred inside the centrifugal devices (Pall Corporation, NY, US) and centrifuged for 10 min, 17000 rcf at 40 °C. After the centrifugation, the samples were transferred into the HPLC vials. 70% of MilliQ water and 30% ACN were used under the isocratic mobile phase. Samples ran under a 1 mL min^−1^ flow and absorbance was measured at 254 nm. Dex release profile from Lipo was assessed by 1000 kDa Float-A-Lyzer dialysis in PBS under agitation at 37 °C. The encapsulation efficiency of Dex was calculated as the ratio between Dex encapsulated after filtration and Dex encapsulated after extrusion measured by HPLC.

### Cryo-Transmission Electron Microscopy

The structure of Lipo and Lipo-Dex was analyzed using Cryo-Transmission Electron Microscopy (Cryo-TEM) at the Baylor College of Medicine Cryo-Electron Microscopy Core Facility (Houston, TX). The Quantifoil R2/1, Cu 200 mesh Holey Carbon grids were pretreated with a 45 s air-glow discharge to make the carbon surface hydrophilic. Alongside these grids, Quantifoil R2/1 200Cu 4 nm thin carbon grids were also glow discharged for 10 s to the efficacy of the added layer of continuous carbon with binding of the Lipo. Vitrification was performed using a Vitrobot Mark IV (FEI, Hillsboro, OR, US) operated at 18 °C and 100% humidity. Each grid had 3 L of Lipo sample applied to it and was subsequently blotted for 1–3 s before being immediately submerged in liquid ethane. The frozen grids were transferred into a JEOL 3200FS microscope (JEOL) outfitted with a Gatan K2 Summit 4kx4k direct detector (Gatan, Pleasanton, CA, US) and a postcolumn energy filter set to 30 eV. Before imaging, the microscope was carefully aligned to prevent beam-induced aberrations or astigmatism that can negatively impact image quality. Images were collected at magnifications of 15.000× and 30.000× with respective pixel sizes of 2.392 and 1.232 angstroms. Images were collected using an exposure time of 1 s with an approximate dose rate of ≈20e-/Å2/s per image. The thickness of the phospholipid bilayer was measured for 10 liposomes using the Fiji software.

### *In Vitro* Studies in Neuronal Cell Culture

SH-Sy5y human and N2a (Neuro-2a) murine cell lines were obtained from Sigma (US) and ATCC (US), respectively. The cells were cultured with Dulbecco’s modified Eagle’s medium (DMEM, Thermo Fisher Scientific, Inc., Waltham, MA, US) supplemented with 10% fetal bovine serum (FBS, Thermo Fisher Scientific) and 1% penicillin-streptomycin. *In vitro* biocompatibility of Lipo-Dex was determined using WST-1 assay. The cells were plated at a density of 5000 cells per well of two 96-well plate and incubated overnight. Cells were treated with Lipo-Dex at the concentration of 0.001, 0.01, 0.1, 1 and 10 µM and incubated for 24 h and 48 h (n=5). WST-1 assay reagent was added after 24 and 48 h incubation for 4 h and absorbance was measured at 490 nm. To access the ability of Lipo-Dex in preventing inflammation, *in vitro,* Sy5y and N2a cells were seeded on a 24-well plate at 1x10^5^ cells per well 24 h prior to the experiments. The cells were pre-treated for 1 h with Lipo-Dex (at Dex concentration of 1 µM and 5 µM) resuspended in a fresh media for 1 h and stimulated with LPS (5 µg/ml) for 24 h (Figure 1b). Cell culture media from each well was collected and stored at -20 °C until analysis. Subsequent cytokine measurements interleukin-6 (IL-6) and tumor necrosis factor-alpha (TNF-α) were done using Milliplex magnetic bead panel (Millipore Sigma, MA, US) according to the manufacturer’s protocol. Standard curves and cytokine concentrations of the samples were generated with the Luminex 200 software.

### Mice and Traumatic Brain Injury Model

Adult male and female (12 weeks old) C57BL/6J mice (Jackson Laboratories, Bar Harbor, ME, US) were housed at the Houston Methodist Research Institute animal facilities. The protocols for all animal studies were reviewed and approved by the Institutional Animal Care and Use Committee (IACUC) at Houston Methodist Research Institute. The mice had ad libitum access to food and water and were kept under a 12 h light and dark cycle. Before and during surgery, mice were anesthetized with isoflurane (3% for induction, 1.5-2% for maintenance). An electromagnetically controlled cortical impact (CCI) injury tool (Impact One stereotaxic impactor, Leica Microsystems, Buffalo Grove, IL, US) was used to perform a left sided moderate TBI at the primary motor and somatosensory cortices as we previously described ^38, 39^ (Figure 1c). The impact site was located at 2 mm lateral and 2 mm posterior to bregma, with a 3 mm diameter flat impact tip, an impact velocity of 3.25 m/s, and an impact depth of 1.5 mm, as previously documented in our laboratory to induce an adequate acute inflammatory response post-injury ^38, 39, 53^. Sham mice underwent all procedures, including anesthesia, but did not undergo cortical impact.

### Biodistribution of Lipo-Dex using IVIS Imaging

Lipo-Dex were labeled with Cy5.5-DSPE and their fluorescence intensity was measured as described above. The labeled Lipo-Dex (100 μL/per mouse) were then administered intravenously via the retro-orbital route into male and female C57BL/6J mice 1 h after CCI injury (n=6 mice per group, 3 males and 3 females). IVIS imaging was performed on the mice, and they were euthanized 24 h after the injury. Then, the mice were euthanized, and their organs (brain, heart, lungs, liver, spleen, and kidneys) and blood were collected and imaged using IVIS to evaluate LNP biodistribution. Serum separation was performed by centrifugation at 4000 rpm for 20 min at 4°C for inflammatory cytokine analysis. The IVIS imaging was conducted using the following acquisition parameters: Ex = 640 nm, Em = 720 nm, Epi-illumination, Bin:(HR)4, FOV:18.4, f2, 0.5 s. The Living Image software was used to analyze the data.

### Cresyl Violet Staining and Lesion Volume Measurements

The brain tissues were fixed in 4% paraformaldehyde overnight, followed by storage in 30% sucrose solution at 4 °C. The brains were then sectioned at 15 μm thickness in the coronal plane through the dorsal hippocampus using a cryostat (Epredia Cryostar NX50, Fisher Scientific, Waltham, MA, US). These sections were then cryoprotected in an antifreeze solution (30% glycerol, 30% ethylene glycol, 40% 0.01M PBS) for storage at -20 °C. To stain the sections, a cresyl-violet solution was prepared in the hood by 0.1% cresyl-violet (Sigma-Aldrich, St. Louis, MO, US) in distilled water, which was then heated while stirred and filtered. The sections were mounted on gelatin-coated glass slides (SuperFrost Plus, ThermoFisher Scientific, IL, US), rehydrated, and stained with the cresyl-violet solution for 10 min. After staining, the slides were dehydrated sequentially with ethanol and cleared in xylene before being covered with Permount (ThermoFisher Scientific) mounting media and coverslipped. The lesion area was assessed on 8 to 12 selected brain sections spaced equidistantly (brain sections at 250 μm intervals) apart, approximately -1.70 to -2.70 mm from bregma. The area of each of the corresponding ipsilateral hemispheres was similarly determined. The lesion volume was obtained by multiplication of the sum of the lesion areas by the distance between sections. The percent lesion volume was calculated by dividing the lesion volume by the total ipsilateral hemisphere volume (similarly obtained by multiplying the sum of the areas of the ipsilateral hemispheres by the distance between sections).

### Organs Paraffin Embedding and Hematoxylin and Eosin Staining

Heart, lungs, liver, spleen, and kidneys were sampled, fixed in 4 % paraformaldehyde for 48 h and transferred to 70 % ethanol. Tissues were processed in a Shandon Exelsion ES Tissue Processor and embedded in paraffin on a Shandon HistoCenter Embedding System, using the manufacturer’s standard processing and embedding protocols. Slides were sectioned at 5 µm thickness. The tissues were dehydrated in 95% ethanol twice for 30 min, followed by soaking in xylene for 1 h at 60-70 °C and paraffin for 12 h. For mouse spleen, liver, and kidneys, 0.5 mL of 95% ethanol was used in the dehydration process. After dehydration, the tissues were stained with hematoxylin solution for 6 h at a temperature of 60-70 °C. Following this, the tissues were rinsed in tap water until the water became colorless. Differentiation of the tissue was carried out using 10% acetic acid and 85% ethanol in water, repeated twice for 2 min, and rinsed with tap water.

### Immunofluorescence Analysis and Cell Death Assay

For immunostaining, parallel brain sections were first washed with PBS-Triton X-100 (0.5%) (PBS-T) for 5 min. The sections were then blocked with 5% normal goat serum (NGS, #1000, Vector Laboratories, Burlingame, CA) for 1 h. Next, brain sections were incubated at 4 °C overnight in PBS-T and 3% of NGS using the following primary antibodies: anti-rabbit Iba-1 (1:500, Wako) for activated microglia/macrophages, anti-rat F4/80 (1:200, R&D Systems) for infiltrated macrophages, anti-rabbit glial fibrillary acidic protein (GFAP) (1:1000, Dako), and anti-mouse S100β (1:200, Sigma). After washing the sections with PBS-T, the corresponding conjugated IgG secondary antibodies, including anti-rabbit or anti-mouse Alexa Fluor 568 and anti-rat Alexa Fluor 488 (all 1:1000, Thermo Fisher Scientific), were added and incubated for 2 h at room temperature. Cell nuclei were counterstained with a DAPI solution diluted in PBS (1:50,000, Sigma-Aldrich), and slides were coverslipped using Tris Buffer mounting medium (Electron Microscopy Sections, Hatfield, PA). To assess cell death, brain sections were processed for DNA strand breaks using Terminal deoxynucleotidyl transferase dUTP nick end labeling (TUNEL) from Fluorescence *In Situ* Cell Death Detection kit (Roche Diagnostic, Indianapolis, IN, US) according to the manufacturer’s instructions.

### Fluorescent *In situ* Hybridization with Immunohistochemical Labeling

Coronal brain sections were mounted on gelatin-coated glass slides (Superfrost Plus, Thermo Fisher Scientific) and stored at −80 °C until use. Fluorescent in situ hybridization (FISH) was performed as per the manufacturer’s instructions using RNAscope® Technology 2.0 Red Fluorescent kit (Advanced Cell Diagnostics (ACD), Hayward, CA, US) as we previously described ^39, 54^. Brain tissue sections were dehydrated by 50%, 70%, and 100% ethanol gradually for 5 min; boiled for 10 min with pretreatment 2 solution (citrate buffer), then incubated with pretreatment 3 solution (protease buffer) for 30 min before hybridization. Sections were incubated at 40 °C for 2 h with the following target probes for mouse: Mus musculus Il1b mRNA (Cat. No. 316891, ACD) and separately with Mus musculus TNF-α (Cat. No.311081, ACD). In addition, the negative (Cat. No. 310043, ACD) and positive (Cat. No. 313911, ACD) control probes were applied and allowed to hybridize for 2 h at 40 °C. The amplification steps were performed according to manufacturer’s directions. The amplification steps were performed according to manufacturer’s directions. After FISH, the slides were washed three times with PBST and blocked with PBST and 5% normal goat serum for 1 h. Immunofluorescence was performed by incubating with polyclonal anti-rabbit Iba-1 (1:500, 019-9741, Wako Chemicals, Richmond, VA, US) overnight at 4 °C for microglia/macrophages cells. Alexa Fluor 488-conjugated goat anti-rabbit (1:1.000, Invitrogen, Carlsbad, CA, US) was applied for 2 h at room temperature. Sections were rinsed with PBS three times and incubated in PBS with DAPI solution for counterstained nuclei. The sections were washed with distilled water and coverslipped with Fluoro-Gel with Tris Buffer mounting medium (Electron Microscopy Sciences).

### Image Analysis of Tissue Damage and Confocal Microscopy Imaging

Images were acquired on a Nikon motorized fluorescence microscope (Eclipse Ni-U, Melville, NY, US) with a pco.edge sCMOS camera (4.2LT USB3) and analyzed using NIS-Elements software. For quantitative analysis of immunolabeled sections, we implemented unbiased, standardized sampling techniques to measure tissue areas corresponding to the injured cortex showing positive immunoreactivity, as we previously described ^25^. Quantitative image analysis of the immunoreactive areas for Iba-1 and GFAP were performed on 15 cortical corpus callosum and hippocampal sections through the level of impact site (AP −2.0 mm) taken with the x20 objective and using the same densitometric analysis method as we previously described ^38, 39^. To quantify the number of Iba-1, S100β, and mRNA-positive cells in the injured cortex, an average of four coronal sections from the lesion epicenter (−1.34 to −2.30 mm from bregma) were counted and imaged for each animal (*n* = 5 mice/group). Within each brain region, every positive cell was counted in each of five cortical fields (x20, 151.894 mm^2^) around the impact area, as we previously described ^39^. Co-localization of IL1-β and TNF-α expression with microglia/macrophages markers was evaluated with *z*-stack acquisitions using the confocal microscope Leica DMi8.

### Rotarod Test

To evaluate motor performance and coordination in the mice, we used a rotating rod apparatus (Ugo Basile Harvard Apparatus, PA, US). Mice were placed on the rod and the speed was gradually increased from 4 to 40 rpm, and the duration of time that the mice were able to stay on the rod was recorded as the latency to fall in seconds. Mice were trained 2 days before CCI injury, and their baseline was established by testing them on the same day of surgery. One day after CCI, mice were tested again (*n* = 5 mice/group).

### Statistical Analysis

The statistical probabilities for size, PDI, ZP, and NPs concentration during formulation steps among different NPs formulations were determined using unpaired t-tests. Meanwhile, for biodistribution studies, statistical probabilities were determined using two-way ANOVA followed by Tukey’s multiple comparison test. For rotarod test and histochemical/immunofluorescence analysis, a two-way ANOVA was used, with time after injury and sex as the variables. A post hoc test with Bonferroni multiple test correction was then applied. All data in the study were presented as mean ± SEM. *p < 0.05, **p < 0.01, ***p < 0.001, and ****p < 0.0001 were considered statistically significant. GraphPad Prism 8 Software (GraphPad; San Diego, CA, US) was used for statistical analysis.

## Acknowledgements

This work was supported by grants from the National Institute for Neurological Disorders and Stroke (NINDS), R21 NS127265 (S.V., B.G.) and TIRR Foundation through Mission Connect grants (SV). G.B. received funding from the Alliance of International Science Organizations (ANSO) Scholarship for Young Talents, University of Chinese Academy of Sciences, College of Material Science, and Opto-electronic Technology. The authors thank the Baylor College of Medicine Cryo-TEM Core Facility for electronic imaging support and the Pathology and Histology Core (HTAP). Figure 1 was created using Biorender.com.

## Author Contributions

G.B., A.Z., F.T., B.G. and S.V. initiated, designed, planned, and oversaw all aspects of the study. G.B., A.Z., H.F., M.H., A.T., S.S., B.G., and S.V. performed the experimental work and data analysis, and they drafted the manuscript. All authors reviewed and edited the final version of the manuscript.

## FIGURE LEGENDS

**Supplemental Figure 1. Lipo and Lipo-Dex toxicity assessment in filtering organs.** Tissue sections of spleen, kidney, lung, and liver underwent Hematoxylin and Eosin (H&E) staining 24 h post-TBI. No obvious tissue differences were observed when comparing the Lipo administrated groups to the Lipo-Dex treated groups. Scale bar represents 50μm.

