## Supplementary figures and images for "Sex-dependent improvement in traumatic brain injury outcomes after liposomal delivery of dexamethasone in mice"

### Supplemental FIgure 1

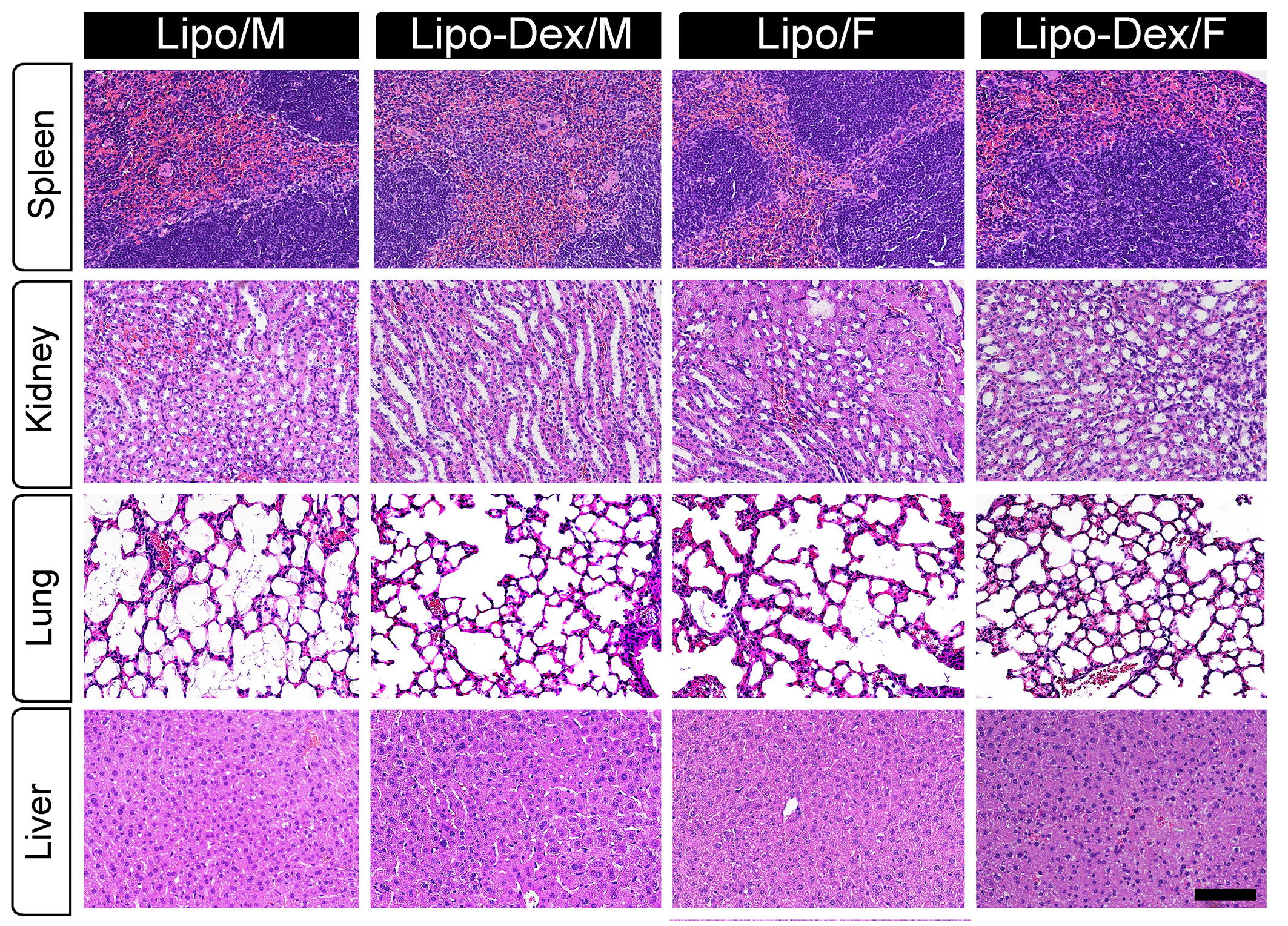
